# In vitro virus-like particle assembly follows two distinct, competing pathways

**DOI:** 10.64898/2026.06.18.733309

**Authors:** Xiaoyan Wang, Hong Luo, Sha Liu, Lukas Gerstweiler

## Abstract

Virus-like particles (VLPs) formed from the murine polyomavirus major capsid protein VP1 are widely used as vaccine antigens and are being explored as nucleic acid and drug delivery vehicles. However, the factors controlling distinct VP1 capsid morphologies remain unclear. We investigated in vitro VP1 assembly with tRNA across NaCl concentrations of 0.15–1.0 M and tRNA mass ratios of 1:1–1:80 (w/w) using SEC-HPLC, transmission electron microscopy, and dynamic light scattering.

Two competing assembly pathways were identified. At low ionic strength (≤0.15 M NaCl), nucleic acid-templated assembly produced compact, tRNA-filled T=1 VLPs (∼28–30 nm). Assembly was maximal at tRNA ratios of 1:10–1:20, whereas excess or insufficient tRNA reduced yields. Increasing NaCl to 0.30 M lowered T=1 yields by 68–97%, and no T=1 particles were detected at ≥0.5 M NaCl.

Conversely, high ionic strength (≥0.5 M NaCl) promoted template-independent formation of hollow T=7 VLPs (∼55–60 nm). T=7 assembly was inhibited by tRNA and was highest without nucleic acid. At 1.0 M NaCl, reducing the tRNA ratio from 1:40 to 1:80 increased T=7 yield more than 11-fold, while removing tRNA produced the greatest assembly efficiency. Kinetic analyses further showed that VP1 concentration and ionic strength regulate nucleation and assembly rate in the template-driven pathway.

These findings show that electrostatic interactions govern pathway selection between tRNA-templated T=1 assembly and salt-driven T=7 self-assembly, providing practical guidance for controlling capsid morphology and cargo loading in VLP-based applications.

## 2. Introduction

Virus-like particles (VLPs) are assembled from viral structural proteins and recapitulate the size and morphology of native virions[1]. The major capsid protein VP1 of polyomaviruses and the related L1 capsid protein from papillomaviruses can self-assemble into VLPs when expressed alone[2, 3]; these VLPs lack genetic material yet retain key structural and antigenic features similar to the source virus[4–6].

Polyomavirus VLPs have been developed across three principal application domains. First, in vaccines: polyomavirus VLPs elicit strong immune responses and serve as prophylactic or therapeutic candidates. Examples include BK and JC polyomavirus VLP vaccines intended to prevent virus reactivation–associated nephropathy or encephalopathy in transplant recipients[7]; immunization with VLPs prevents primary virus infection and restrains growth of virus-associated tumors and has been advanced as a tumor immunotherapy platform, such as HER2/neu-expressing tumors.[8–10] Second, as gene and drug delivery systems: polyomavirus VLPs self-assemble while packaging exogenous nucleic acids, can function as vectors for gene transfer or as drug carriers. For instance, SV40 VLPs reassembled with pGL3 (5.3 kb) exhibit efficient hepatic transduction following hydrodynamic tail-vein injection in rats or mice, and reassembly with the suicide gene PE38 results in pronounced antitumor effects in xenograft models[4, 11, 12]. Third, in diagnostics and serology: VP1 pentamers or VP1-based VLPs act as capture antigens in enzyme immunoassay, and because VLPs preserve conformational epitopes, VLP-based enzyme immuno-assay (EIA) is typically more sensitive than assays using monomeric VP1 antigens and enables competitive inhibition with heterologous VLPs to assess cross-reactivity[13].

Two operational assembly strategies are described for polyomavirus VP1: high salt induced, and template induced. In low-salt or physiological buffers, charged surface regions on VP1 experience electrostatic repulsion that prevent assembly[14, 15]. Elevated salt screens surface charges, reducing repulsion, and increasing hydrophobic interactions and promoting tight pentamer–pentamer association and assembly[16]. Under 1 M NaCl or 0.5 M (NH4)_2_SO_4_ at pH 7.2, purified VP1 pentamers form native capsid-like structure with sedimentation coefficients of approximately 120–300 S (peak around 144 S, comparable to native empty capsids), and electron microscopy shows capsid-like particles together with pentameric units[14].

In template-induced assembly, the positive N-terminal region of VP1 (lysine- and arginine-rich) binds the negatively charged nucleic acid backbone. In the presence of DNA or RNA, VP1 adsorbs electrostatically onto the nucleic acid, multiple pentamers align along the same strand, and the nucleic acid acts as a template that nucleates pentamer clustering and drives closure into a shell[17, 18]. In DNA-driven systems, near-physiological ionic strength (0.15 M NaCl) with a small amount of divalent cation to stabilize inter-capsomer contacts (2 mM CaCl₂) supports formation of well-ordered particles across pH 5–9. E.g. a 4729 bp circular DNA template, sub stoichiometric-to-stoichiometric ranges (DNA:VP1 pentamer molar ratios of approximately 1:144 to 1:72) favour 40 nm, T=7-like particles[19]. In RNA-driven systems, short flexible single-stranded RNA (approximately 500 nt) serves as an intrinsically bendable scaffold. In assembly buffers containing 250 mM NaCl and 100 mM MOPS at pH 7.2, cooperation with VP1 pentamers yields homogeneous T=1 icosahedral particles containing one RNA and 12 pentamers per particle[20].

For mechanistic studies, Schmidt et al. showed that the spontaneous *in vitro* assembly of the major capsid protein VP1 from murine polyomavirus is a reversible equilibrium that can be stabilized by oxidizing inter subunit disulfide bonds, which prevent dissociation and are not strictly required for assembly but are important for obtaining complete and stable particles[21]. Garcea et al. reported that truncation of the polyomavirus C terminal domain blocks the transition from pentameric capsomeres to complete capsids[22]. Nilsson et al., using cryo electron microscopy, reconstructed BK polyomavirus VLPs and found two symmetry classes, T=1 and T=7, governed by the conformational flexibility of the C-terminal arms and tunable linkages at local threefold axes[23]. These findings highlight the heterogeneity of VLP assembly, with the same capsid protein capable of forming structurally distinct particles. A comprehensive, unified theoretical framework for VLP assembly has yet to be established.

## 3. Methods

### VP1 Expression

The wild type of murine polyomavirus VP1 gene (GenBank: M34958.1) was cloned into the pETDuet-1 expression vector and transformed into E. coli Rosetta™ 2 (DE3) Singles™ cells (Merck, Germany). Transformation was carried out via heat shock at 42 °C for 30 seconds, followed by recovery in S.O.C. medium (0.5% yeast extract (Thermo Fisher Scientific, P0021), 2% tryptone (Thermo Fisher Scientific, LP0042), 10 mM Sodium Chloride (NaCl) (Chem Supply, SL046), 2.5 mM KCl, 10 mM Magnesium Chloride Hexahydrate (MgCl₂·6H₂O) (Chem Supply, MA029), 10 mM Magnesium Sulphate (MgSO_4_) (Chem Supply, ML073), 20 mM glucose (Chem Supply, GA018)) and plating on Terrific Broth (TB) agar (1.2% tryptone, 2.4% yeast extract, 0.4% glycerol (Chem Supply, GA010), and 89 mM phosphate buffer (17 mM Potassium Dihydrogen Orthophosphate Anhydrous (KH₂PO₄) (Chem Supply, PA009) and 72 mM di-Potassium Hydrogen Orthophosphate Anhydrous (K₂HPO₄) (Chem Supply, PA020)), supplemented with 100 µg mL⁻¹ ampicillin (Chem Supply, GA0283) and 35 µg mL⁻¹ chloramphenicol (Chem Supply, GA0258). A master cell bank was established by culturing a single colony in TB medium containing the same antibiotics and storing the resulting cultures at −80 °C in 25% v/v glycerol. For protein expression, overnight seed cultures were inoculated into 300 ml fresh TB medium with antibiotics and incubated in 1 L Erlenmeyer shake flask with shaking at 200 rpm (at 37 °C) until the optical density at 600 nm (OD₆₀₀) reached 0.8–1.0. Protein expression was induced by adding 0.1 mM Isopropyl β-D-1-thiogalactopyranoside (IPTG) (Thermo Fisher Scientific, 15529019), followed by incubation at 27 °C for 16 h. Cells were harvested by centrifugation at 6000 × g for 10 min at 4 °C (Beckman Coulter centrifuge, model JPG21B002). The cell pellets were washed once with saline (0.9% w/v NaCl), aliquoted, and stored at −80 °C for further use.

For cell lysis, the pellets were resuspended in a Tris-based buffer (20 mM Tris (hydroxymethyl) aminomethane (Tris) (Chem Supply, GB4431), 30 mM 1,4 Dithiothreitol (DTT) (Chem Supply, DL131), 10 mM Ethylenediaminetetraacetic Acid Disodium salt dihydrate (EDTA) (Chem Supply, EA023), 5% glycerol, pH 8.9) and disrupted by ultrasonication on ice (Scientz JY92-IIDN) at 460 W using 3 s on / 5 s off cycles for 35 min. The lysates were clarified by centrifugation at 15,000 × g for 40 minutes at 4°C.

### VP1 Purification

VP1 capsomere purification followed a protocol recently published by our group[24]. All operations were performed on an ÄKTA™ Advance system (Cytiva, 2639144) using a 12 mL self-packed Capto™ MMC column. Clarified lysate from the previous step was adjusted to approximately 0.35 M NaCl to match the MMC loading conditions and then loaded directly onto the Capto™ MMC column for purification. The MMC loading buffer was 20 mM Tris, 5 mM DTT, 2 mM EDTA, 5% w/w glycerol, and 0.35 M NaCl, pH 8.9, which promoted strong binding of VP1 on MMC while suppressing DNA/RNA – VP1 interactions[25, 26]. Clarified lysates were loaded at 3 mL min⁻¹. After loading, the column was washed with 5 column volumes (CV) of loading buffer at 5 mL min⁻¹ to remove unbound proteins. Elution was performed by a stepwise shift to alkaline conditions using an elution buffer composed of 20 mM Di-Sodium hydrogen phosphate (Chem Supply, SO03391000), 5 mM DTT, 2 mM EDTA, and 5% w/w glycerol, pH 11.8, at 3 mL/min, yielding the main capsomere peak. Elution was monitored at A280 and A260. Fractions then were collected, aliquoted and stored at −80°C.

### VP1 polishing

Frozen aliquots were thawed and filtered through 0.22 µm filter (Filter-Bio, cat. no. FBS25MCE022S). Then the VP1 polishing was performed on an ÄKTA™ Advance system equipped with a Superose® 6 Increase 10/300 GL column at a flow rate of 0.6 mL·min⁻¹. The column was equilibrated with SEC buffer (20 mM Tris-HCl, 5 mM DTT, 2 mM EDTA, 5% w/w glycerol, 0.5 M NaCl, pH 8.0), with an equilibration volume of at least 1.2 CV. Elution was carried out isocratically over 1.5 CV, and the capsomere containing peak collected. The protein and potential nucleic acid carryover were monitored by A280 and A260UV signal[25–27].

### Bradford Assay

Bradford working solution was prepared by mixing 1 mL Bradford reagent (Bio-Rad, cat. no. 5000001) with 4 mL Milli-Q water to a final volume of 5 mL. A bovine serum albumin (BSA) standard curve spanning 0.2 to 0.9 mg/mL was prepared.

Standards and samples were mixed with the working solution at 1:50 (v/v; sample to working-solution ratio), protected from light, and incubated for at least 5 min at room temperature. Absorbance at 595 nm was measured in triplicate using a microplate reader (SPECTROstar Nano, 8000-86). VP1 concentrations were determined by interpolating sample absorbance values from the BSA standard curve.

### Salt-driven VLP assembly

Purified VP1 capsomeres were adjusted to the desired protein concentration in 20 mM Tris-HCl, pH 7.0, with various NaCl concentration. For assembly at different NaCl concentration the SEC eluate (already at 0.5 M NaCl) was first adjusted to the desired NaCl concentration adding concentrated Tris-HCl/NaCl (3 M NaCl), then diluted with the corresponding buffer (e.g 20 mM Tris-HCl, 1.0 M NaCl (pH 7.0) to the desired protein concentration (e.g. 0.3 mg/mL). Samples were briefly mixed by flipping followed by microcentrifugation (OHAUS FC5306; 5 s) and incubated at 25 °C for 12–24 h to allow spontaneous VLP assembly. As a reference, VLP assembly was carried out in assembly buffer containing 20 mM Tris-HCl (pH 7.0) and the indicated NaCl concentration according to the standard procedure described in our previous publication[26].

### tRNA-templated VLP assembly

This procedure requires a tRNA template, purified VP1 capsomeres, and Tris HCl buffers with defined NaCl concentrations. Yeast tRNA (75 nt; 10 mg/mL stock; Thermo Fisher Scientific, cat. no. 2989577) served as the assembly template. VP1 capsomeres obtained from the preceding purification in 0.5 M NaCl with DTT at the concentration of 1 to 1.5 mg/mL.

Unless otherwise indicated, reactions were adjusted to 0.30 mg/mL VP1. Tris–HCl buffers containing 0–3.0 M NaCl were prepared in advance to adjust the final NaCl concentration and the desired tRNA: VP1 mass ratios.

To achieve the lower NaCl concentration than at the final mixture, the required volume of Tris–HCl (pH 7.0) without NaCl to offset salt contributed by the VP1 stock buffer and the tRNA solution were calculated and added. The mixtures were then adjusted to achieve the final concentration of 0.30 mg/mL VP1 by adding Tris–HCl containing the target NaCl concentration for each condition.

For assembly setup, 0.1 to 10 µL of tRNA stock (prepared in 0 M NaCl) was dispensed into a 1.5 mL microcentrifuge tube and kept on ice. The pre-determined Tris-HCl buffer at the intended NaCl concentration (0, 0.15, 0.2, 0.3, 0.4, 0.5 or 1 M) was added and gently premixed with the tRNA to avoid premature diluting VP1, which can trigger spontaneous self-assembly. Purified VP1 was added last, and the mixture was gently homogenized by pipetting to minimize shear induced aggregation. All steps were performed on ice. Alternatively, VP1 and tRNA were added together. Please note, buffer should not add directly to VP1 prior to introducing tRNA.

The mixture was briefly consolidated in a mini centrifuge for 5 s, then incubated at room temperature for 12 to 24 h with the tube cap loosened to promote DTT oxidation and allow disulfide bond formation between capsomeres. A 500 mL beaker was placed inverted over the tubes to limit dust contamination and reduce evaporation. Caps were then closed and the beaker removed, followed by an additional 12 to 24 h incubation at room temperature. During this period, the tubes were gently tilted (30 to 45 °) once or twice to ensure homogeneity.

### Size Exclusion-High-Performance Liquid Chromatography (SEC-HPLC)

All reactions were analysed by SEC-HPLC using a Shimadzu UFLC-XR system (LC-20AD-XR pump, SIL-20AXR autosampler, SPD-M20A diode array detector, CTO-20 column oven). Separations used a Superose 6 Increase 10/300 GL column (Cytiva) with Tris–HCl eluent (pH 7.2, 0.5 M NaCl). UV was monitored at 260 and 280 nm. The injection volume was 50 μL for all experiments. The total run time was 180 min, the flow rate was 0.25 ml/min, and the column temperature was maintained at 25 °C.

### Dynamic Light scattering (DLS)

DLS was performed using a Zetasizer Nano ZS (Malvern Panalytical, MAL1289521). Samples were diluted one to five with Milli Q water to maintain the scattering intensity within the instrument’s recommended range and to minimize multiple scattering. Disposable plastic cuvettes (BRAND, 7590-15) were used. The instrument software was set to 25 °C with an equilibration period of 120 s and at least three replicate measurements per sample.

### Transmission electron microscopy (TEM)

Samples (5 μl) were diluted 1:10 (v/v) with Milli-Q water and applied onto carbon-coated copper grids (ProSciTec, GSCU100C). After incubation for 5 min, excess liquid was removed. Negative staining was performed with 2% (w/v) Uranyl Acetate Dihydrate, EM stain (UO₂(OCOCH₃)₂·2H₂O) (ProSciTech, C079) for 2 min. Imaging was conducted on an FEI Tecnai G2 Spirit transmission electron microscope equipped with an Olympus SIS Veleta CCD camera at 120 kV.

### Statistical analysis

All quantitative data were presented as the mean ± standard deviation (SD). Statistical analyses were performed in GraphPad Prism 11. Data are presented as mean ± SD. Significance levels were set at p < 0.05, p < 0.01, and p < 0.001. Values with p < 0.05 were considered statistically significant.

## 4. Results

### 4.1 Salt concentration selects assembly pathway between T=1 and T=7 VLP

To determine whether NaCl concentration governs the assembly of T=1 or T=7 VLPs*, in vitro* reactions were performed by mixing tRNA with VP1 pentamers at mass ratios from 1:1 (w/w) to 1:80 (w/w). NaCl was tested across 0.15–1.0 M. Assembly products were analysed by SEC-HPLC. Peaks corresponding to T=1 and T=7 VLPs (Fig. S1c, d) were integrated, and the areas were normalized to permit quantitative comparison across conditions.

#### Low NaCl concentration promotes RNA-mediated T=1 VLP assembly

The highest yield of T=1 VLPs was measured at 0.15 M NaCl, across all tRNA:VP1 mass ratios from 1:1 (w/w) to 1:80 (w/w) (Fig. 1). Increased NaCl concentrations of 0.30 M markedly suppressed T=1 VLP assembly. For example, at tRNA:VP1 = 1:10 (w/w) T=1 VLP yield decreased by 67.97% by increasing NaCl from 0.15 M to 0.3 M (Fig. 1b). The difference was supported by t-test (t = 4.99, df = 3.87, P = 0.0082; mean difference = 0.68; 95% CI, 0.30 to 1.06). The same trend held at tRNA:VP1 = 1:20 (w/w) (decrease of 93.22%) (Fig. 1c; 0.15 M: 1.00 ± 0.07 vs 0.30 M: 0.07 ± 0.005; *t* = 13.32, df = 2.02, *p* = 0.0054; mean difference 0.93; 95% CI 0.63–1.23) and tRNA:VP1 = 1:40 (w/w) (decrease of 97.19%) (0.15 M: 1.00 ± 0.07 vs 0.30 M: 0.028 ± 0.003; *t* = 13.06, df = 2.01, *p* = 0.0058; mean difference 0.97; 95% CI 0.65–1.29). No T=1 peak above baseline (baseline + 3 SD) was detected at 0.50 or 1.00 M for any ratio, indicating complete suppression of T=1 assembly at higher ionic strength.

**Fig 1.**
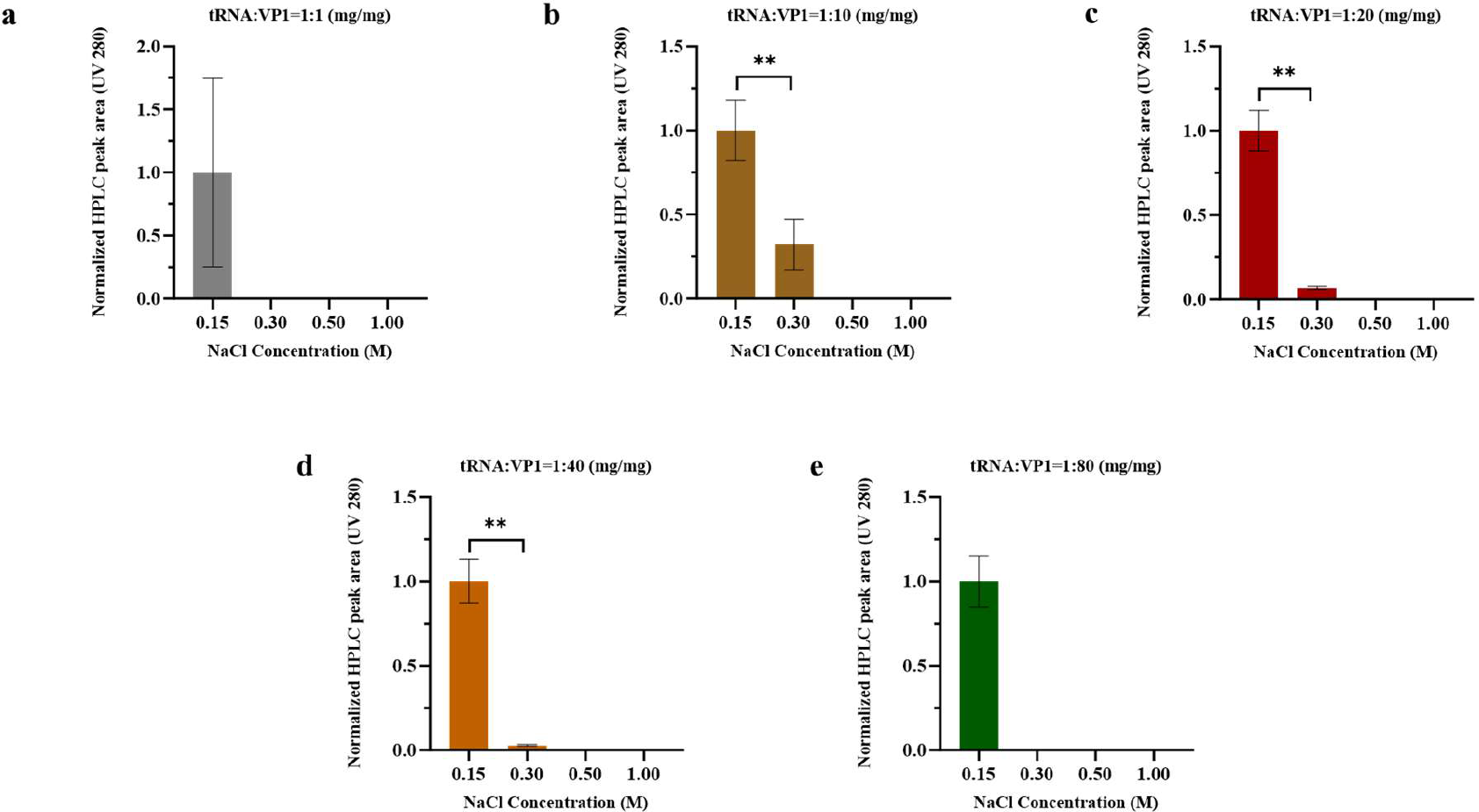
HPLC-SEC quantification of the assembled VLP–tRNA peak, with peak area integrated and normalized. tRNA: VP1 mass ratios: (a) 1:1 (w/w), (b) 1:10 (w/w), (c) 1:20 (w/w), (d) 1:40 (w/w), (e) 1:80 (w/w). Yields of T=1 VLP–tRNA were compared at 0.15, 0.3, 0.5, and 1.0 M NaCl. Error bars represent ± SD (n = 3). Statistical analysis used t-test with the following significance rule: ns, p ≥ 0.05; * p < 0.05; ** p < 0.01; *** p < 0.001.

#### High NaCl concentration drives tRNA-independent assembly of T=7 VLPs

In contrast, T=7 VLP assembly occurs at higher NaCl with lower ratio of tRNA template. At tRNA:VP1 = 1:40–1:80 (w/w), no discernible T=7 peak was observed at both 0.15 and 0.30 M NaCl concentration, whereas increasing NaCl to at least 0.50 M started to produce a detectable T=7 VLP peak with 1:40 (w/w) ratio (Fig. 2a). The T=7 peak area reached the maximum at 1.0 M NaCl (43.78% higher than at 0.5 M NaCl). The similar trend was seen at 1:80 (w/w), where the observed T=7 VLP yield at 1.00 M NaCl was 13.8 fold higher than that at 0.5 M NaCl.

**Fig. 2.**
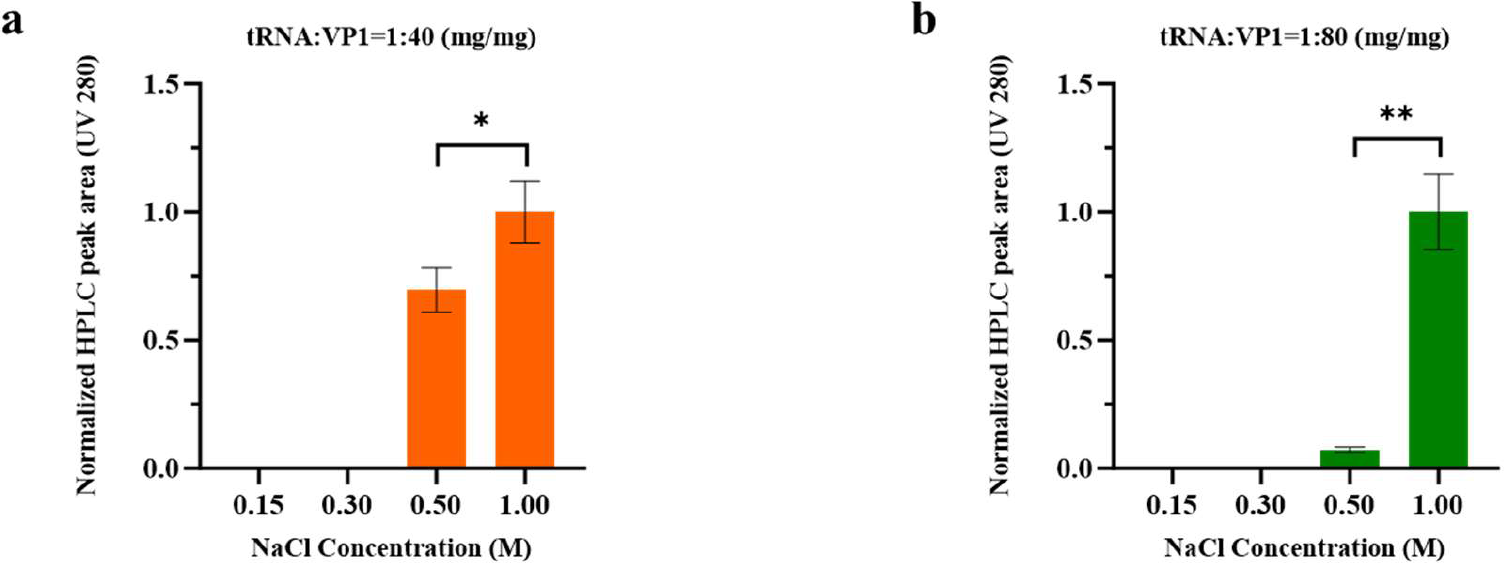
Influence of NaCl on assembly of T=7 VLPs. Assembly was assessed by HPLC-SEC; the T=7 VLP elution peak was quantified by SEC with peak areas integrated and normalized. Tested tRNA: VP1 mass ratios: (a) 1:40 (w/w), (b) 1:80 Yields were compared at 0.15, 0.3, 0.5, and 1.0 M NaCl. Higher ratios of tRNA to capsomere did not lead to formation of T=7 VLPs (presented in Fig. 6. Error bars represent ± SD (n = 3). Statistical evaluation applied t tests with significance denoted as: ns, p ≥ 0.05; * p < 0.05; ** p < 0.01; *** p < 0.001.

#### Structural validation of NaCl-dependent VLP morphologies by TEM

To confirm the morphology of T=7 VLPs assembled at elevated NaCl conditions, TEM was performed to visualize the products generated either without tRNA or with tRNA across multiple salt conditions. Four tRNA groups were examined: 0.5 M NaCl at tRNA:VP1 mass ratios of 1:40 (w/w) and 1:80 (w/w), and 1.0 M NaCl at 1:40 (w/w) and 1:80 (w/w). Two controls were included: 1.0 M NaCl with VP1 only and VP1 with standard assembly buffer from literature[24]. At 0.5 M NaCl, VP1 pentamers predominantly formed incompletely or mis formed shells, with a minor fraction of spherical particles comparable in size to canonical T=7 VLPs. This pattern is most evident in the TEM micrographs for 0.5 M NaCl at a tRNA:VP1 mass ratio of 1:80 (w/w) (Fig. 3a and 3b). The findings indicate that the SEC-HPLC peak attributed to T=7 VLPs at 0.5 M NaCl might correspond to incompletely assembled particles. Raising NaCl to 1 M yielded distinct T=7 VLPs whose diameters matched those of native polyomavirus T=7 particles assembled in high-salt assembly buffer without tRNA. The corresponding SEC-HPLC elution positions were likewise similar to those of VLPs assembled without tRNA (Fig. 3b, 3c, 3f; Fig. S1 b). TEM micrographs showed hollow interiors across VLPs, indicating no encapsulated tRNA. Consistently, SEC-HPLC at 1 M NaCl with a tRNA:VP1 mass ratio of 1:80 (w/w) yielded a UV260 to UV280 peak height ratio of 0.86 at the T=7 VLP elution, consistent with nearly empty particles (Fig. 3b, c; Fig. S1 c). To further verify that tRNA did not participate in T=7 assembly, VLPs were generated at 1 M NaCl without tRNA; TEM still showed T=7 particles formed by VP1 pentamers with size and morphology indistinguishable from those obtained in the presence of tRNA (Fig. 3e).

**Fig. 3.**
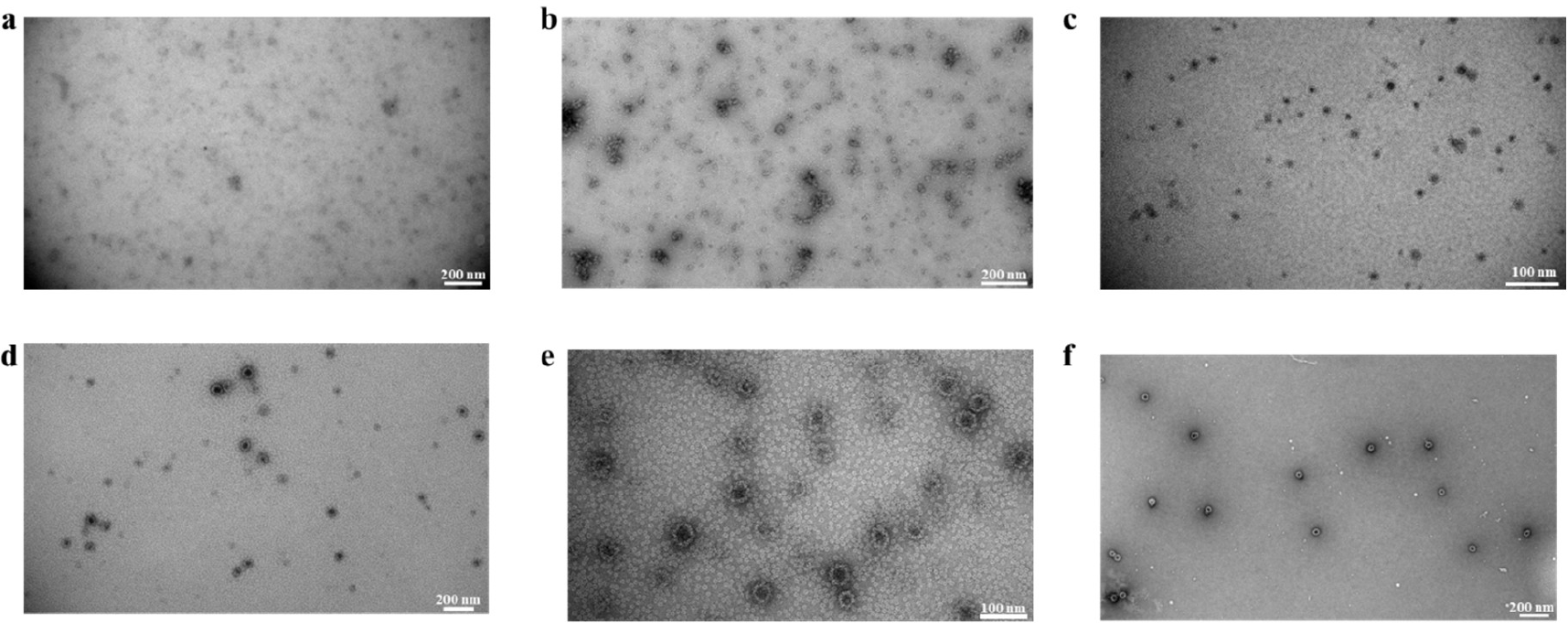
TEM of VP1 assemblies formed with or without tRNA under varying NaCl concentrations. (a) 0.5 M NaCl, tRNA:VP1 mass ratio 1:40 (w/w); (b) 0.5 M NaCl, 1:80 (w/w); (c) 1.0 M NaCl, 1:40 (w/w); (d) 1.0 M NaCl, 1:80 (w/w); (e) 1.0 M NaCl, VP1 assembly without tRNA; (f) VP1 assembly in Assembly Buffer.

#### DLS confirms particle homogeneity and assembly completeness

TEM results indicated that, irrespective of tRNA, high NaCl concentrations induced VP1 to assemble into T=7 spherical VLPs. To further quantify and compare the particle size distribution and assembly homogeneity in solution, samples were analysed by DLS. Under 0.5 M NaCl with tRNA present (tRNA:VP1 mass ratio 1:80), three DLS peaks were detected: 14 nm corresponding to unassembled VP1 pentamers, 204.9 nm representing partially closed or mis-formed VLP assembly intermediates consistent with TEM observations, and 1699 nm indicating aggregation (Fig. 4a). At 0.5 M NaCl without tRNA, a single peak at 115.4 nm was detected, with intensity markedly higher than the peaks corresponding to empty VLPs and to tRNA loaded VLPs (Fig. S1a; Fig. 4d), indicating that 0.5 M NaCl alone did not yield fully closed T=7 VLPs. At 1 M NaCl with a tRNA:VP1 mass ratio of 1:80, particles formed with a mean diameter of 56.38 nm (Fig. 4c). A secondary peak on the left of the main peak averaged 12.85 nm, consistent with the size of unassembled VP1 pentamers. The main peak closely matched that of empty VLPs at 54.54 nm (Fig. S1 a). VP1 assembly was likewise evaluated without tRNA. Under this condition, the main peak averaged 59.83 nm, and a second peak at 4469 nm indicated aggregation (Fig. 4d).

**Fig. 4.**
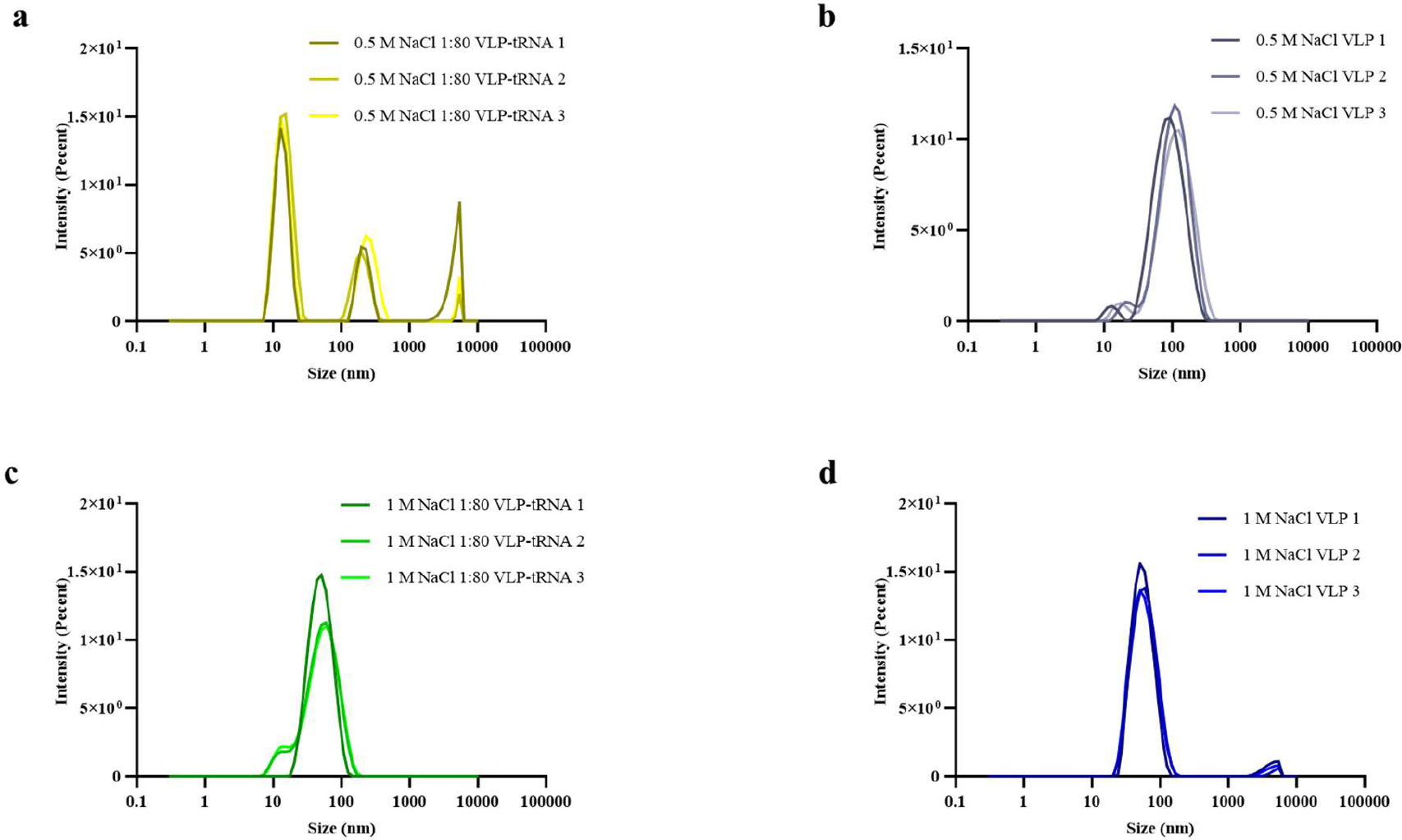
Dynamic light scattering (DLS) analysis of VP1 assembly behaviour under varying NaCl concentrations with or without tRNA. (a) 0.5 M NaCl, tRNA:VP1 = 1:80; (b) 0.5 M NaCl, without tRNA; (c) 1.0 M NaCl, tRNA:VP1 = 1:80; (d) 1.0 M NaCl, without tRNA. Peaks correspond to VP1 pentamers, partially assembled intermediates, and aggregates as indicated.

### 4.2 Influence of RNA:VP1 ratio on assembly pathway

To assess the effect of the tRNA:VP1 mass ratio on the T=1 and T=7 assembly pathway, reactions were prepared at NaCl concentrations of 0.15, 0.30, 0.50, and 1.00 M with tRNA:VP1 ratios of 1:1 (w/w), 1:10 (w/w), 1:20 (w/w), 1:40 (w/w), and 1:80 (w/w). Assembly efficiency was quantified by integrating and normalizing the SEC–HPLC UV 280 nm peak areas corresponding to T=1 and T=7 VLPs. At 0.50 and 1.00 M NaCl, no T=1 VLP signal above baseline was detected at any ratio, so these conditions were excluded from subsequent analyses.

#### RNA:VP1 ratio controls T=1 VLP assembly efficiency

At 0.15 M NaCl (Fig. 5a), a tRNA:VP1 ratio of 1:10 (w/w) produced the largest T=1 peak area (normalized to 1.00 ± 0.12, n = 3). The 1:20 (w/w) condition was comparable (normalized 0.94 ± 0.10, n=3; P > 0.05 vs 1:10), whereas 1:1 (w/w) gave a near-baseline signal (≈0) but showed aggregation (peak at excluded column volume). Reducing the tRNA fraction beyond 1:20 (w/w) reduced assembly, with peak areas reduced by 70.20% (normalized 0.30 ± 0.05 (n=3)) at 1:40 (w/w) and reduced by 90.90% (normalized 0.09 ± 0.02 (n=3) at 1:80; both were significantly lower than 1:10 (w/w) and 1:20 (w/w) (P < 0.01, t test). These data indicate that a tRNA:VP1 ratio between 1:10 (w/w) and 1:20 (w/w) is optimal for T=1 assembly under low ionic strength, while higher tRNA concentrations lead to aggregation.

**Fig. 5.**
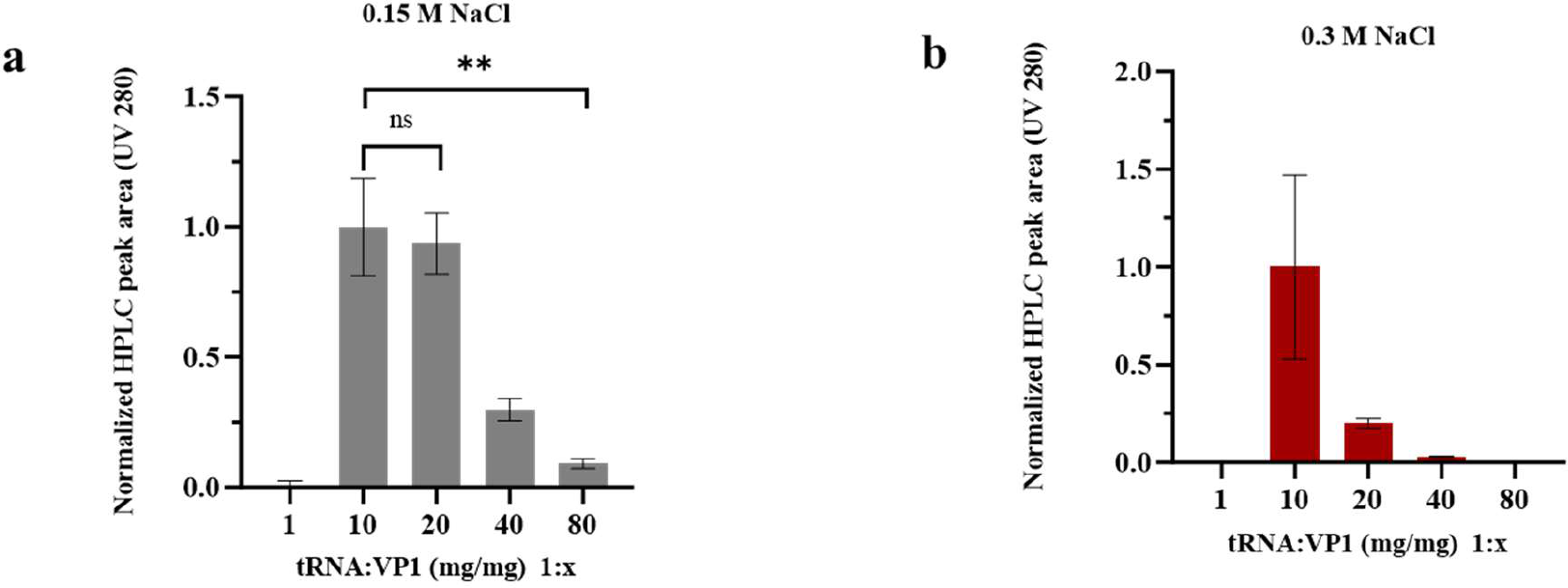
Effect of tRNA: VP1 mass ratio and NaCl concentration on T=1 VLP assembly efficiency quantified by SEC–HPLC. (a) At 0.15 M NaCl, the normalized HPLC peak area for T=1 VLPs was maximal at a tRNA:VP1 ratio of 1:10 (w/w), indicating optimal template availability for nucleation. Assembly efficiency decreased sharply at higher (1:40 (w/w), 1:80 (w/w)) or lower (1:1 (w/w)) ratios, consistent with inhibition under RNA excess or template deficiency. (b) At 0.3 M NaCl, a similar trend was observed, with the 1:10 ratio yielding the highest peak area. Excess tRNA (1:1 (w/w)) suppressed assembly, while insufficient template (1:40–1:80 (w/w)) markedly reduced T=1 VLP formation. Error bars represent ± SD (n = 3). Statistical evaluation applied t tests with significance denoted as: ns, p ≥ 0.05; * p < 0.05; ** p < 0.01; *** p < 0.001.

At 0.30 M NaCl (Fig. 5b), no T=1 peak was observed at a 1:1 (w/w) tRNA:VP1 ratio (but aggregated peak in exclusion volume of SEC column). Similar to 0.15 M NaCl the 1:10 (w/w) ratio yielded the highest signal. Decreasing the ratio to 1:20 (w/w) reduced the normalized peak area to 0.20 ± 0.02 (n=3), and at 1:40 (w/w) and 1:80 (w/w) the T=1 peaks were scarcely detectable (normalized area < 0.05). These results indicate that at moderate ionic strength, efficient T=1 assembly requires an intermediate tRNA:VP1 ratio, while both excess tRNA can lead to aggregation and insufficiency of tRNA suppress VLP formation.

#### RNA suppresses T=7 VLP formation

To assess the effect of the tRNA on assembly of T=7 VLPs under salt induced assembly, SEC–HPLC was applied to reactions containing 0.50 or 1.0 M NaCl with tRNA:VP1 ratios from 1:1 (w/w) to no tRNA. Assembly efficiency was quantified from normalized UV 280 nm peak areas assigned to T=7 VLPs. Data at 0.15 and 0.30 M NaCl were not analyzed further, since no T=7 peak exceeding baseline plus 3 SD was detected under these lower salt conditions.

At 0.50 M NaCl (Fig. 6a), high tRNA:VP1 ratios from 1:1 (w/w) to 1:20 (w/w) produced no detectable T=7 peak. Reducing the tRNA fraction to 1:40 (w/w) generated a detectable T=7 signal, with a normalized peak area of 0.315 ± 0.039 (n = 3). A further reduction to 1:80 (w/w) produced a comparable signal of 0.364 ± 0.052 (n = 3). The 1:40 (w/w) and 1:80 (w/w) groups did not differ significantly (P = 0.27, t test). In the absence of tRNA, the T=7 peak area increased further and was set as the no-template control, with a normalized value of 1.000 ± 0.416 (n = 3). These results indicate that excess tRNA suppresses salt-driven T=7 assembly at 0.50 M NaCl, whereas low or no tRNA levels allow partial T=7 assembly.

**Fig. 6.**
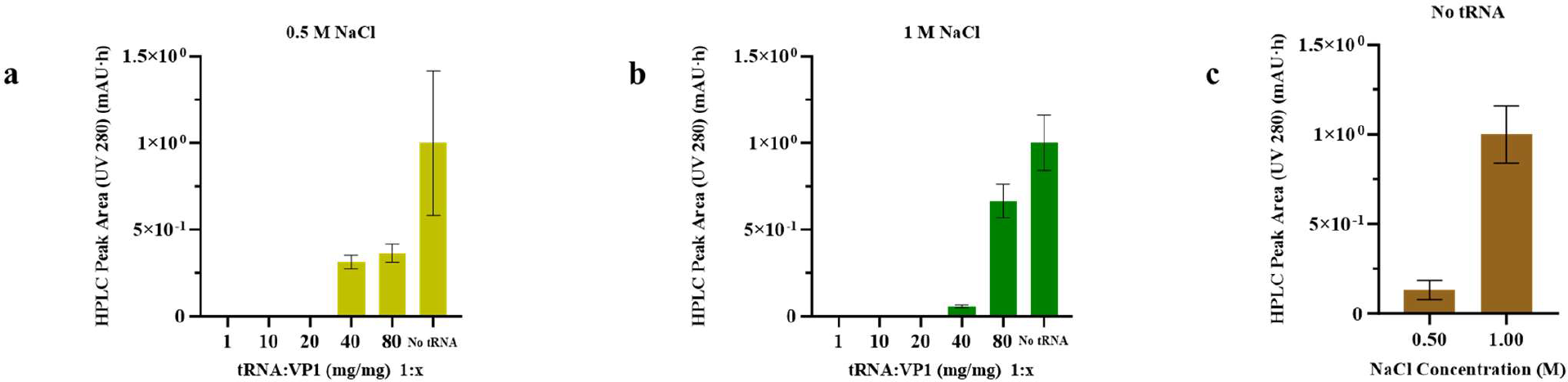
Effect of tRNA: VP1 ratio on T=7 VLP assembly under high NaCl conditions. (a) At 0.5 M NaCl, no detectable T=7 peak was observed at tRNA: VP1 ratios from 1:1 to 1:20 (w/w). Partial assembly emerged at 1:40 and 1:80, yielding normalized peak areas of 0.315 ± 0.039 and 0.364 ± 0.052, respectively, while the no-tRNA control exhibited the highest signal (1.000 ± 0.416). (b) At 1.0 M NaCl, T=7 peaks were absent at ratios from 1:1 to 1:20; however, peak intensity increased from 0.060 ± 0.007 at 1:40 to 0.665 ± 0.096 at 1:80. The no-tRNA control again showed the highest signal (1.000 ± 0.159). (c) Comparison of no-tRNA conditions at 0.5 M and 1.0 M NaCl, with normalization to the 1.0 M group, revealed enhanced T=7 peak areas at higher salt concentration, indicating that NaCl promotes T=7 assembly in the absence of tRNA. Error bars represent mean ± SD (n = 3).

At 1.0 M NaCl (Fig. 6b), the inhibitory effect of tRNA on T=7 assembly was more evident. tRNA:VP1 ratios from 1:1 (w/w) to 1:20 (w/w) produced no detectable T=7 peak (0.00 ± 0.00, n = 3). Lowering the tRNA fraction to 1:40 (w/w) generated only a weak T=7 signal, with a normalized peak area of 0.060 ± 0.007 (n = 3). A further reduction to 1:80 (w/w) markedly increased the T=7 peak area to 0.665 ± 0.096 (n = 3). The difference between 1:40 (w/w) and 1:80 (w/w) was significant (P < 0.01, independent samples t test). In the absence of tRNA, the T=7 peak area increased further and was set as the no-template control, with a normalized value of 1.000 ± 0.159 (n = 3). These data show that high ionic strength induces T=7 VLP assembly and that tRNA interferes with this process, as higher tRNA concentrations effectively suppress salt driven assembly and low tRNA concentrations increase yield.

To further assess NaCl-dependent assembly of tRNA-free T=7 VLPs, template-free samples prepared at 0.50 M and 1.0 M NaCl were compared (Fig. 6c). After normalization to the mean peak area of the 1.0 M NaCl group, the T=7 peak area was 0.132 ± 0.055 at 0.50 M NaCl and 1.000 ± 0.159 at 1.0 M NaCl, corresponding to an approximately 7.6-fold increase. These results indicate that, in the absence of tRNA, increasing the NaCl concentration from 0.50 M to 1.0 M enhances salt-driven assembly of T=7 VLPs.

#### Structural validation of tRNA dosage effects on VLP-tRNA morphology by TEM

To verify the morphology of T=1 VLPs assembled at different tRNA: VP1 ratios, TEM was performed on reactions prepared at mass ratios of 1:1 (w/w), 1:10 (w/w), and 1:20 (w/w). At 1:1 (w/w), VP1 pentamers formed irregular aggregates with no discernible higher order capsids (Fig. 7a). At 1:10 (w/w) (Fig. 7b), numerous uniformly dispersed spherical particles of approximately 30 nm in diameter were observed; particle edges were well defined, and the interiors showed homogeneous electron density consistent with a filled core. This morphology differed from that of T=7 VLPs, which display a hollow interior, and suggests incorporation of tRNA within the T=1 particles. In agreement with this interpretation, the corresponding T=1 SEC–HPLC peak at this ratio exhibited a UV260/UV280 ratio of approximately 1.20, indicative of nucleic acid containing protein particles (Fig. S1d).

**Fig. 7.**
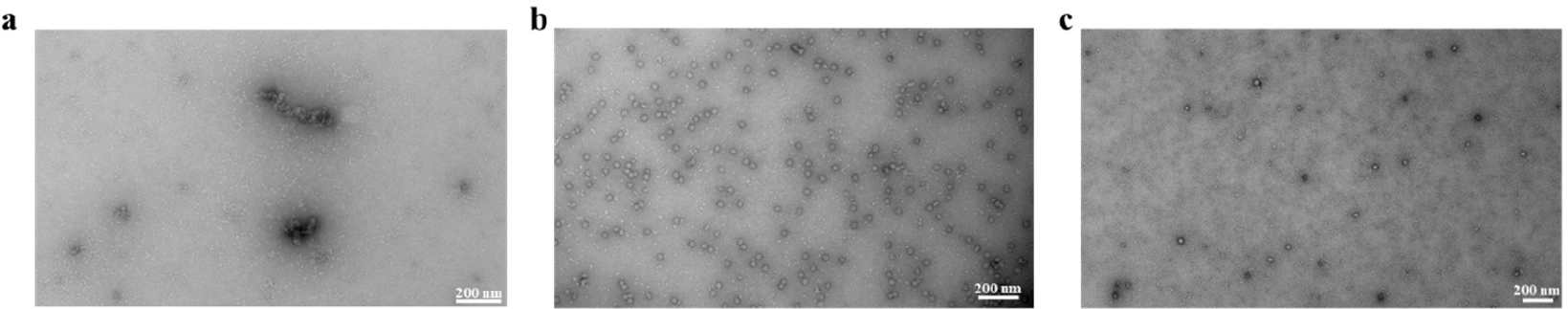
TEM analysis of tRNA-dependent T=1 VLP assembly at 0.15 M NaCl. Transmission electron micrographs show the effect of tRNA:VP1 mass ratio on T=1 VLP formation. (a) At tRNA: VP1 = 1:1 (w/w), only irregular VP1 aggregates were observed, with no ordered structures. (b) At 1:10 (w/w), abundant, well-dispersed spherical T=1 VLPs (∼30 nm) with solid cores and uniform density were formed, indicating efficient assembly with incorporated tRNA. (c) At 1:20 (w/w), VLPs form T=1 with solid core. Scale bars, 200 nm.

At a tRNA:VP1 ratio of 1:20 (w/w) (Fig. 7c), T=1 VLPs retained intact virus-like morphology with a solid core. Collectively, these data indicate that VP1 pentamers use tRNA as a template to nucleate assembly and produce structurally complete, well dispersed T=1 VLPs at intermediate tRNA:VP1 ratios, whereas excess tRNA suppresses T=1 assembly and yields disordered VP1 aggregates, consistent with the SEC-HPLC profiles showing a peak in the excluded volume.

#### DLS confirms RNA-dependent control of VLP-tRNA size and homogeneity

To quantify particle size distributions and assess the effect of tRNA dosage on T=1 VLP assembly, DLS was conducted at tRNA:VP1 mass ratios of 1:1 (w/w), 1:10 (w/w), and 1:80 (w/w). At 1:1 (w/w), two principal peaks were observed: a 15.2 nm species consistent with VP1 in solution and a 1200 nm peak indicative of aggregation, with no peak at VLP size, in agreement with TEM and SEC (Fig. 8a). At 1:10 (w/w), a single dominant peak at 27.64 nm (PDI 0.128) was detected, matching the expected size of T=1 VLPs (Fig. 8b; Fig. S1a). At 1:80 (w/w) (Fig. 8c), a dominant peak at 20.12 nm, approximately 3.3 nm larger than unassembled VP1, was observed, which is likely an overlay between the unassembled capsomeres and the low yield T=1 VLPs (Fig. 5a). DLS also showed an additional 276.8 nm peak corresponding to irregular aggregates; These data are consistent with SEC-HPLC results, showing that low levels of tRNA (1:80) yield low levels of T=1 VLPs, whereas excess tRNA (1:1 (w/w)) promotes aggregation; an intermediate ratio of 1:10 (w/w) supports the formation of stable, uniform T=1 VLPs.

**Fig. 8.**
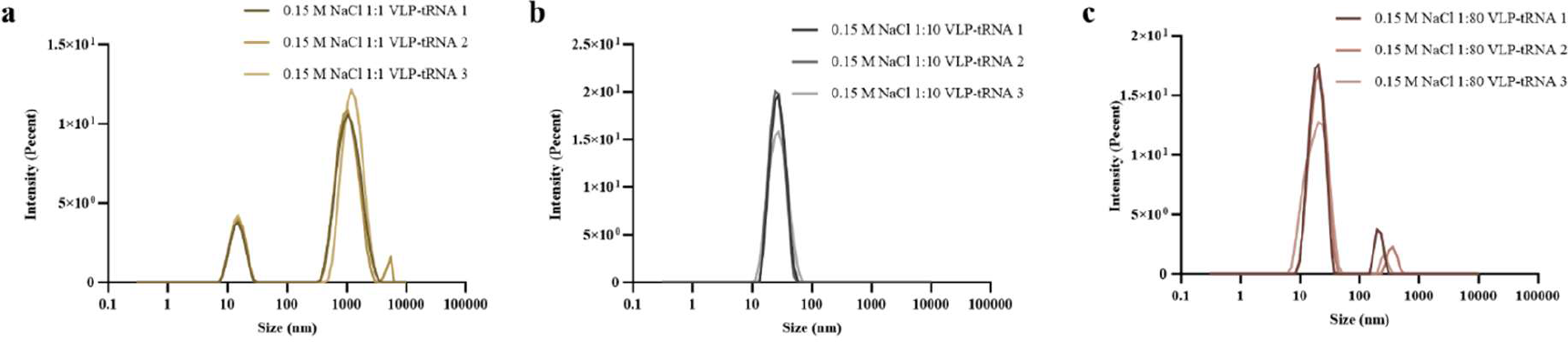
Dynamic light scattering (DLS) characterization of tRNA-dependent T=1 VLP assembly at 0.15 M NaCl. Particle size distributions at different tRNA: VP1 mass ratios reveal the effect of template concentration on assembly. (a) At 1:1 (w/w), only VP1 capsomere (∼15 nm) and aggregation (∼1200 nm) peaks were detected, with no VLP formation. (b) At 1:10 (w/w), a dominant peak at ∼28 nm (PDI = 0.128) corresponded to uniform T=1 VLPs. (c) At 1:80 (w/w), peaks at ∼20 nm and ∼277 nm indicated small oligomers and aggregates, confirming insufficient template for VLP assembly.

### 4.3 Kinetic study on template assembly pathway

Screening identified a tRNA:VP1 mass ratio of 1:10 (w/w) that maximized tRNA encapsulation, and this ratio was used in subsequent SEC–HPLC analyses to quantify the kinetics of the template guided assembly pathway.

#### VP1 concentration modulates the efficiency of template-driven assembly

To assess the effect of VP1 concentration under low NaCl conditions, VP1 was tested at 0.1, 0.2, and 0.3 mg/mL (Fig. 9a). Reducing VP1 from 0.3 to 0.2 mg/mL decreased the VLP–tRNA peak area at 48 h by 51.0%, and a further reduction to 0.1 mg/mL produced an additional 16.9% decrease. This shows that the assembly yield of T=1 VLPs is not only dictated by the tRNA ratio but also by the VP1 concentration.

**Fig. 9.**
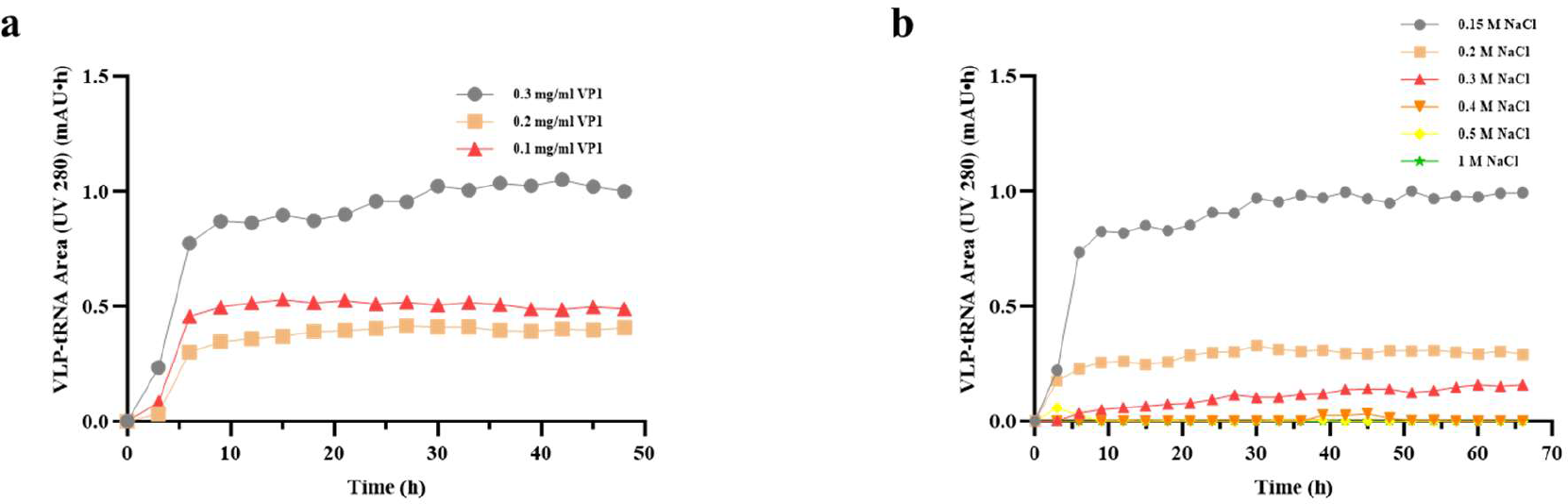
Kinetic analysis of tRNA-driven VP1 assembly under varying protein and salt concentrations. (a) Effect of VP1 concentration (0.1–0.3 mg/mL) on tRNA encapsulation efficiency at 0.15 M NaCl and a tRNA:VP1 ratio of 1:10 (w/w). Higher VP1 concentrations accelerated assembly. (b) Effect of NaCl concentration (0.15–1 M) on tRNA-mediated VLP formation at 0.3 mg/mL VP1. Efficient assembly occurred below 0.3 M NaCl, while higher ionic strength (≥0.5 M) suppressed tRNA-VLP formation.

#### NaCl concentration influences the kinetics and extent of template-driven assembly

The effect of NaCl concentration on the kinetics of template-driven assembly was then examined by varying NaCl concentration (Fig. 9b). At NaCl concentrations ≥ 0.5 M, no VLP–tRNA peak was detected at 66 h. While lowering NaCl concentration generally increases T=1 VLP yield. At 0.4 M NaCl, a small VLP–tRNA peak (area 0.032) appeared between 39 and 48 h and then gradually declined to zero. At 0.3 M NaCl, the peak area increased more slowly, reaching 0.16 (normalized to the 0.15 M NaCl condition at 66 h) at 66 h. In contrast, at 0.15 M NaCl, assembly proceeded most rapidly during the first 6 h, with the VLP–tRNA peak area rising from 0 to 0.73, which is 146.2% higher than the 66 h peak area observed at 0.2 M NaCl. From 6 to 30 h, the assembly rate approached a steady state, and from 30 to 66 h the reaction gradually reached a plateau, reaching maximum at 66 h. Overall, the 66 h peak area at 0.15 M NaCl exceeded that at 0.2 M NaCl by 240.8%, indicating that both low ionic strength and sufficient VP1 concentration are critical for efficient template-driven VLP assembly.

## 5. Discussion

Using polyomavirus VP1 with a 75 nt tRNA template, this study shows that low NaCl conditions and a tRNA:VP1 mass ratio of 1:10 to 1:20 (w/w) favour the formation of T=1 VLPs composed of 12 pentamers that encapsulate tRNA. Evidence includes the solid particle morphology observed by TEM and an elution peak in SEC with a 260:280 ratio greater than 1. In contrast under high NaCl conditions, even low levels of tRNA, e,g, VP1:tRNA mass ratios of 1:80 (w/w) compared to no tRNA reduce the assembly yield of T=7 VLPs comprising 72 pentamers, which appear hollow in TEM images. The schematic summarizes a dual pathway for polyomavirus VP1 capsid assembly, the experimental data support a model in which VP1–nucleic acid interactions influence pathway selection. (Fig.10).

**Fig. 10.**
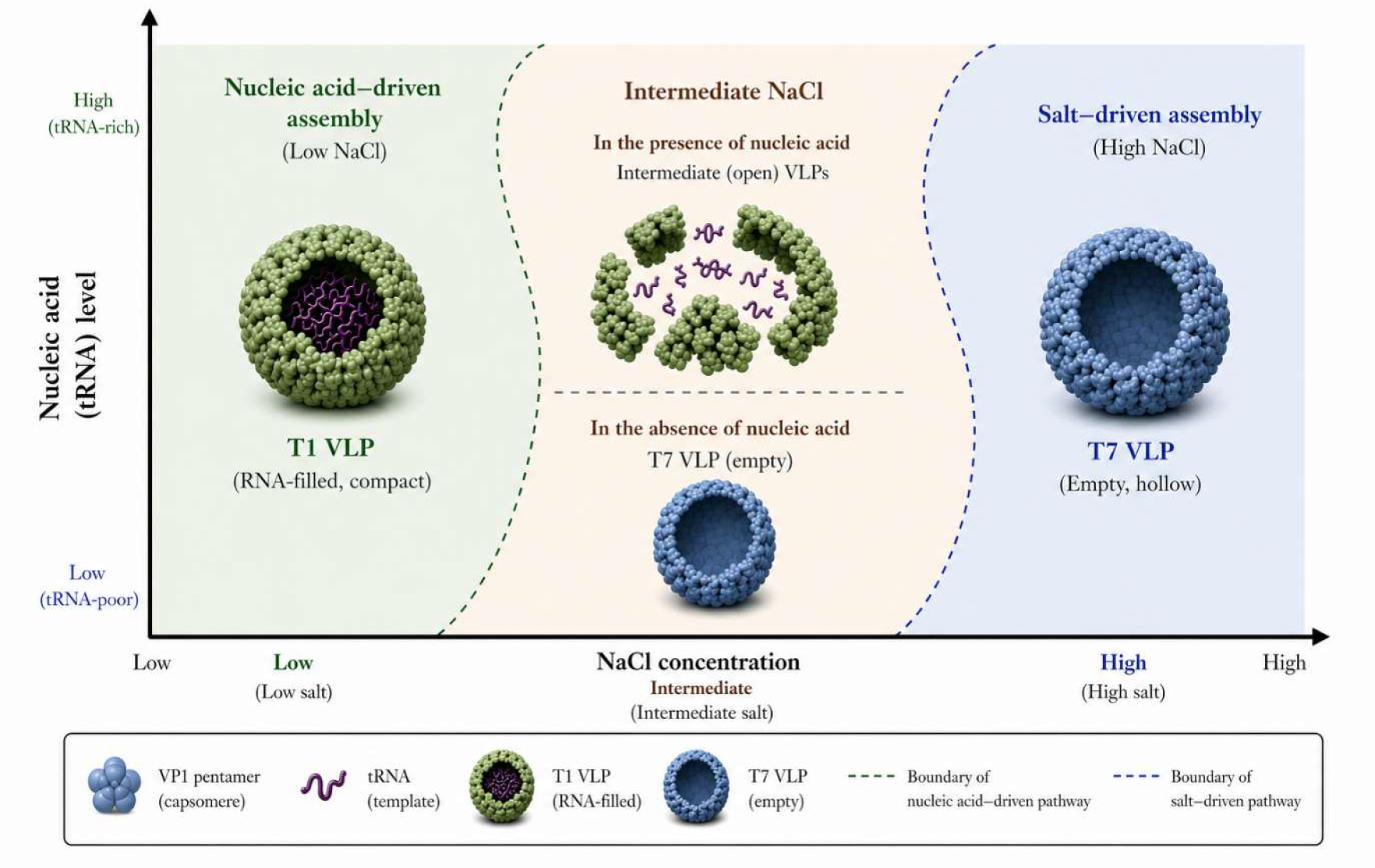
Distinct assembly pathways of Polyomavirus VP1. At low NaCl concentration, assembly is nucleic acid driven, producing compact T=1 VLPs that encapsulate tRNA, with lower NaCl and higher tRNA typically increasing yield. At high NaCl concentration, assembly follows a salt driven pathway and yields hollow T=7 VLPs, with higher NaCl and low levels of tRNA increasing yield. At intermediate NaCl concentration, VP1 tends to assemble into intermediate structures in the presence of nucleic acid, whereas hollow T=7 VLPs are formed in the absence of nucleic acid.

### 5.1 Effect of NaCl concentration on VLP assembly pathway selection and efficiency

#### T=1 VLP

At tRNA:VP1 mass ratios from 1:1 (w/w) to 1:80 (w/w), increasing NaCl concentration progressively reduced the yield of T=1 tRNA loaded VLPs, and at 0.3 M NaCl formation of loaded T=1 VLPs was nearly abolished. A plausible explanation is that negatively charged tRNA promotes local enrichment of positively charged VP1 pentamers through electrostatic interactions, which may facilitate nucleation[19]. NaCl ions shield these interactions, weakening the attraction between nucleic acid and protein and suppressing the template driven assembly pathway. At lower salt concentrations (e.g. 0.15 M NaCl), electrostatic attraction is stronger, which favours VP1 association with tRNA and ordered assembly; at higher NaCl, stronger ionic screening lowers the electrostatic interaction between VP1 and tRNA, slows the assembly rate, and yields fewer loaded particles. These trends align with the template driven assembly hypothesis for VP1, in which nucleic acids promote oriented association of capsomers through electrostatic attraction[19, 20]. Under high salt conditions (>0.3 M NaCl), ionic screening may weaken VP1–tRNA interactions and reduce the ability of tRNA to mediate productive nucleation. Similar behavior has been reported in polyomavirus VP1 systems, where increased ionic strength favors asymmetric or polymorphic aggregates rather than ordered T=1 structures[14, 15, 28]. Gerstweiler et al. (2021) further showed that the presence of nucleic acids and modulating electrostatic interactions at high salt concentration can promote disordered aggregation of protein nucleic acid complexes, thereby blocking the productive assembly pathway[20]. The observed trend of high salt suppressing loading indicates that higher ionic strength weakens the template adsorption driving force between VP1 and tRNA, which reduces loading efficiency. These findings accord with the concept of assembly governed by electrostatic balance proposed by Garcea et al. (1986), indicating that VP1 assembly is highly sensitive to the ionic environment and that moderate electrostatic attraction is essential for formation of regular T=1 VLP[14].

#### T=7 VLP

As NaCl concentration increased, at a tRNA:VP1 mass ratio of 1:40 (w/w) to 1:80 (w/w), VLP morphology shifted markedly. SEC-HPLC indicated that at 0.15 M to 0.3 M NaCl, no proper T=7 VLP particles were formed; a distinct T=7 VLP signal emerged at 0.5 M and reached the highest yield at 1.0 M. TEM showed partially closed T=7 capsids at 0.5 M NaCl. Raising NaCl to 1.0 M produced abundant closed T=7 particles which size and morphology closely matched T=7 capsids formed by spontaneous assembly in the absence of tRNA, both in assembly buffer and at 1.0 M NaCl. DLS corroborated these trends: at 0.15 M the size distribution was broad and unstable; at 0.5 M a dominant peak near 50 nm appeared; at 1.0 M a narrower single peak indicated a homogeneous and stable particle population. It suggests that increasing ionic strength shifts the system from nucleic acid template driven loading toward protein driven assembly, and that the presence of nucleic acids impair the salt driven assembly pathway. Under low salt conditions, tRNA as a negatively charged template can attract positively charged VP1 pentamers through electrostatic interactions, initiating nucleation and favouring loading into small capsids such as T=1[19, 20]. Conversely, Na⁺ and Cl⁻ effectively screen long-range electrostatic attraction, decreasing the binding affinity between tRNA and VP1[29]. Once electrostatic attraction is weakened, tRNA no longer functions as a template and assembly becomes dominated by protein-protein interactions among VP1 pentamers[25, 26]. The process thus transitions from template driven assembly to spontaneous protein assembly that might be driven by hydrophobic interactions. This low salt loading followed by high salt self-assembly behaviour aligns with the energetic basis of VP1 assembly: electrostatic attraction under low salt favours compact nucleic acid loaded small capsids such as T=1, whereas charge screening at high salt both diminishes this attraction and reduces subunit repulsion, rendering the larger T=7 geometry thermodynamically preferred[15, 30]. In addition, the T=7 architecture of polyomavirus capsids depends on the stability of a pore at the fivefold axis, maintained by electrostatic and hydrogen bonding interactions between adjacent VP1 pentamers; ionic strength modulates these interfacial interactions and shifts the balance between incompletely closed intermediates and fully closed capsids[31]. This mechanistic picture is consistent with the presence of unclosed intermediates at 0.5 M NaCl and the predominance of fully closed T=7 particles at 1.0 M.

Overall, increasing NaCl drives a transition from tRNA template driven loading that favours T=1 particles to spontaneous VP1 assembly that yields T=7 capsids. Higher ionic strength likely attenuates electrostatic attraction between VP1 and tRNA, reducing template-mediated loading, while screening repulsive interactions between VP1 pentamers. This charge-screening effect may favour pentamer association by hydrophobic interactions and formation of the larger T=7 architecture[14, 29, 32].

### 5.2 Effect of tRNA template concentration on pathway selection and assembly efficiency of VLPs

#### T=1 VLP

In vitro assembly of T=1 VLPs by polyomavirus VP1 is sensitive to the concentration of the tRNA template. Under 0.15 to 0.30 M NaCl, formation of T=1 tRNA loaded VLPs showed a pronounced dependence on the tRNA:VP1 mass ratio. SEC-HPLC, TEM and DLS consistently showed that efficient T=1 assembly yield was highest near 1:10 to 1:20 (w/w), indicating a narrow stoichiometric window for productive tRNA driven assembly. Collectively, these results delineate a narrow stoichiometric window near 1:10 to 1:20 (w/w) in which the tRNA template supports efficient T=1 assembly under moderate ionic strength. The ratio dependence likely reflects a cooperative balance between electrostatic attraction and subunit contact energy in template driven assembly. At lower ionic strength, the negatively charged tRNA serves as a template that enriches positively charged VP1 pentamers on its surface through electrostatic attraction and induces nucleation, thereby initiating assembly[19, 20]. An intermediate amount of tRNA provides sufficient nucleation sites without over neutralization of charge, yielding maximal assembly within the 1:10 (w/w) to 1:20 (w/w) range. In contrast, when nucleic acid is in excess, for example at 1:1 (w/w), the surplus negative charge promotes formation of nonspecific complexes in which VP1 pentamers bind multiple tRNA molecules, disrupting spatial coordination among subunits and suppressing ordered polymerization[33, 34]. This charge oversaturation effect shifts the system from controlled loading to disordered aggregation, consistent with the DNA saturation dependent nucleation model proposed by Mukherjee et al. (2010)[35]. At low tRNA content, such as 1:40 (w/w) or 1:80 (w/w), the number of template sites becomes insufficient to recruit multiple VP1 pentamers for concurrent binding and nucleation, which slows nucleation and favours unstable oligomers[32]. Under these conditions, VP1 occurs predominantly as non-assembled capsomeres reflected by a minor DLS peak near 20 nm in agreement with TEM. A moderate nucleic acid concentration maintains electrostatic attraction and pentamer contact energy within an optimal window, enabling ordered and efficient T=1 loading. Excess or deficiency of nucleic acid disrupts this energetic balance, producing aggregation or nucleation failure, respectively. These observations are consistent with prior studies of VP1 template driven assembly and reinforce the conclusion that T=1 VLP assembly is governed by template concentration[18–20, 32, 33].

#### T=7 VLP

Under high salt conditions (0.5–1.0 M NaCl), the effect of the tRNA:VP1 ratio on T=7 VLP formation reversed relative to T=1 loading. SEC–HPLC quantification detected no T=7 VLP–associated peaks at high tRNA:VP1 ratios (1:1 (w/w) to 1:20 (w/w)), indicating that excess tRNA markedly suppresses productive assembly of VP1 into T=7 capsids. Reducing the ratio to 1:40 (w/w) and 1:80 (w/w) produced a substantial increase in assembly efficiency, with the complete absence of tRNA showing even higher assembly yield at both 0.5 M and 1.0 M NaCl. These observations suggest that, at elevated ionic strength, even small amounts of tRNA may perturb VP1 self-assembly and reduce the formation of closed T=7 particles. The trend aligns with a shift in the dominant assembly driver from template-induced to spontaneous assembly. Under low salt conditions, VP1 assembly is primarily template driven by nucleic acids; tRNA promotes pentamer enrichment and nucleation through electrostatic attraction. Under high salt, electrostatic screening reduces tRNA–VP1 attraction, diminishing the dominance of the nucleic acid template. The pathway appears to shift toward spontaneous assembly, in which pentamer–pentamer interactions become more dominant[14, 15]. One possible explanation is that excess negatively charged tRNA interacts with multiple VP1 pentamers and perturbs the inter-pentamer organization required for productive T=7 closure. The mechanistic interpretation remains hypothetical, as this effect has not been described in the literature. Relatively, Fuertes et al. (2023)[29] reported that elevated ionic strength reshapes the energy landscape of capsid assembly, lowers repulsive barriers between subunits, and makes attractive interactions between proteins the dominant term. This charge-screening rationale could explain why T=7 self-assembly predominates at high salt and why the presence of a tRNA template becomes inhibitory. Consistent with this view, real time single molecule measurements by van Rosmalen et al. (2020)[32] showed that when binding between the template and protein weakens, VP1 assembly transitions from dependence on nucleic acids to a self-limiting pathway, with the rate proportional to the pentamer association energy.

Accordingly, at elevated ionic strength the dominant determinant of VP1 assembly shifts from electrostatically driven templating to pentamer–pentamer interactions that are largely non-electrostatic. Charged tRNA seems to perturb the self-organisation of the capsomeres reducing the assembly yield. Efficient formation of T=7 VLPs therefore requires complete depletion of nucleic acids, which has not been described as a critical factor yet. The outcome is consistent with early polyomavirus VP1 self-assembly studies[14] and with recent charge regulated assembly models[18, 29, 32], supporting the interpretation that high-salt T=7 assembly is not dependent on nucleic acid templating.

Template concentration showed opposite effects on T=1 and T=7 assembly. Moderate tRNA:VP1 ratios (1:10–1:20 (w/w)) favour the formation of T=1 VLPs, while excessive template (1:1 (w/w)) was associated with nonspecific aggregation, possibly due to charge over-neutralization or nonproductive VP1–tRNA complexation. Low template levels (1:40–1:80 (w/w)) reduced T=1 assembly, which may reflect insufficient template-mediated nucleation. In contrast, T=7 assembly achieves optimal efficiency with no tRNA and already trace amounts (1:80 (w/w)) markedly reduced assembly yield.

### 5.3 Kinetic analysis of the template assembly pathway

Given extensive work by Middelberg et al. on how NaCl influences assembly of polyomavirus VP1 into T=7 VLPs[36–41], the discussion turns to its effects on the formation of T=1 VLPs.

#### Assembly kinetics of T=1 VLP

At a fixed tRNA:VP1 mass ratio of 1:10 (w/w), increasing NaCl concentration markedly reduced the rate of tRNA driven VP1 assembly. At 0.15 M NaCl, assembly progressed rapidly during the first 6 h, followed by a slower growth phase that approached equilibrium. At 0.3 M NaCl and higher, assembly was nearly abolished, indicating that ionic strength is a critical determinant of the assembly kinetics. This trend accords with prior reports. Ionic strength reshapes the assembly energy landscape by screening the electrostatic attraction between positively charged VP1 pentamers and the negatively charged tRNA template[34, 42]. Under low salt conditions, stronger VP1–tRNA electrostatic attraction may lower the effective nucleation barrier, thereby promoting faster particle formation; at higher salt, ion screening weakens template to protein interactions, decreases the nucleation rate, and extends the lag phase[43]. These data are consistent with a model in which the rate of tRNA-induced VP1 assembly is influenced by the balance between electrostatic attraction and ionic screening. Intermediate ionic strength, approximately 0.15 M, maintains sufficient attraction while limiting nonspecific protein aggregation, enabling efficient template directed assembly; at higher NaCl, electrostatic interactions are screened and assembly is impeded, approaching kinetic arrest.

#### Impact of VP1 concentration on T=1 VLP assembly

At a tRNA:VP1 mass ratio of 1:10 (w/w), the tRNA induced assembly rate and yield increased with rising VP1 concentration. This concentration dependence indicates that VP1 exerts a key regulatory effect on the nucleation step of template driven assembly. Low VP1 concentration reduces the probability of productive collisions between pentamers, limiting nucleation events and slowing assembly; increasing VP1 concentration elevates nucleation frequency and accelerates assembly. A possible reason is that VP1 concentration may influence local pentamer enrichment around the tRNA template and thereby affect the probability of productive nucleation. Mukherjee et al. demonstrated that SV40 VP1 assembly on DNA shows a sigmoidal dependence on VP1 concentration, with a Hill coefficient of 5.8 ± 0.6, indicating highly cooperative assembly involving approximately 6–7 VP1 pentamers. This suggests that near the assembly threshold, small reductions in VP1 concentration can disproportionately lower assembly yield. DNA may therefore act not only as a binding substrate, but also as a scaffold that concentrates multiple VP1 pentamers or pentameric clusters on the same molecule, promoting cooperative nucleation[44]. In line with this mechanism, stable spherical particles are formed only at elevated VP1 fractions[35], and single-molecule studies show that low VP1 concentrations favor intermediate accumulation, whereas high concentrations accelerate progression from nucleic acid engagement to closed capsids[32].

## 6. Conclusion

This study delineates two distinct assembly pathways of polyomavirus VP1 that can interfere with each other. Under low-salt conditions, appropriate nucleic acid (here tRNA) VP1 ratios enable nucleation and assembly driven by tRNA as a template, ultimately yielding structurally stable T=1 VLP–tRNA particles. In contrast, under high-salt conditions, the nucleic acid–driven pathway is suppressed even in the presence of tRNA and hollow T=7 VLPs are generated, while increasing levels of tRNA suppress the assembly and reduce yield. These pathways are not independent but competitive; when the salt-driven pathway predominates, excessive tRNA interferes with the formation of T=7 VLPs and can completely suppress assembly at high levels. Likewise, no stable T=1 VLPs are formed at high salt, even at favourable tRNA:VP1 ratios. These findings provide both practical guidance for the large-scale production of VLPs and in-vitro assembled viral vectors and important reference conditions for VLP vaccine design, where precise modulation of salt and nucleic acid concentrations allows controlled generation of either empty or cargo-loaded capsids, and identifies remaining nucleic acids as a key driver for assembly yield and efficiency.

## 7. Author statements

### 7.1 Author contributions

X.W.: Investigation, Formal analysis, Data curation, Visualization, Writing – original draft. H.L.: Methodology, Writing – review & editing. S.L.: Supervision, Writing – review & editing. L.G.: Conceptualization, Methodology, Formal analysis, Supervision, Project administration, Writing – review & editing. All authors reviewed and approved the final version of the manuscript.

### 7.4 Ethical approval

Ethical approval was not required because this study did not involve human participants, human material, human data, or animal experiments.

### 7.5 Consent for publication

Not applicable. This article does not contain any personal details, images, videos, or other information that could identify an individual.

## 7.6 Acknowledgements

The authors acknowledge the technical support and instrumentation access provided by the relevant microscopy and analytical facilities at Adelaide University.

